# Connecting growth and yield models to continuous forest inventory data to better account for uncertainty

**DOI:** 10.1101/2025.03.14.643281

**Authors:** Malcolm S. Itter, Andrew O. Finley, Aaron Weiskittel

## Abstract

Models of forest growth and yield are frequently used to inform adaptive management decisions aimed at increasing forest resilience or promoting long-term carbon storage. Despite the increasing ecological detail represented in growth and yield models, there remains large variability (uncertainty) in predictions of forest dynamics under global change. Quantifying this uncertainty and accounting for it when making management decisions is integral to sustainable management in the face of changing conditions. However, the structure and complexity of modern growth and yield models make it challenging to quantify uncertainty and propagate it to predictions of forest dynamics under alternative management strategies. To address this challenge, we develop a Bayesian dynamical model informed by continuous forest inventory data that supports the quantification and propagation of uncertainty in predictions of forest dynamics at a stand scale. The model predicts the temporal evolution of the size-species distribution using a matrix projection process model approximating growth, mortality, and regeneration. Disturbance is integrated through its effect on the size-species distribution within a stand providing a flexible framework to represent adaptive management. We apply the model to long-term inventory data from the Penobscot Experimental Forest in Maine, USA to predict multi-decadal biomass dynamics under five alternative management strategies. Predictions are used to identify the management strategy maximizing live aboveground biomass growth and yield over the model period. We conclude by discussing the benefits and challenges of connecting the model to large-scale inventory data and how its predictions can be used to better inform adaptive management decisions.

## Introduction

Models of forests dynamics (FDMs) are an important management tool for predicting forest responses to silvicultural treatments. These predictions are used to assess and compare the ecological and economic outcomes of alternative management scenarios under the broad heading of growth and yield modeling (Pretzsch, 2009). Here, growth and yield modeling is used to describe the application of FDMs that explicitly represent management allowing for prediction of forest growth, mortality, and regeneration under specified management strategies (Weiskittel et al., 2011). Given the long time frames over which forest management outcomes are realized (decades to centuries) these predictions are frequently relied upon to make management and policy decisions.

While FDMs have a long history of application in sustainable forest management (Pretzsch, 2009), interest in these models and their predictions of growth and yield has increased rapidly in the current era of global change (Maréchaux et al., 2021). FDMs are often applied to predict adaptive management outcomes under novel climate and disturbance scenarios quantifying the capacity of simulated management to sustain ecosystem services or promote resistance and resilience to changing conditions (Mina et al., 2017; Seidl et al., 2018; Gustafson et al., 2020). Further, with growing interest in forest carbon for climate change mitigation, FDMs are increasingly applied to predict forest carbon outcomes under alternative management approaches (Seidl et al., 2008; Nunery and Keeton, 2010; Ameray et al., 2021). These predictions may be used to quantify the carbon benefits of Improved Forest Management practices aimed at increasing carbon stocks in forests over time as part of broader natural climate solutions (Griscom et al., 2017; Giffen et al., 2022; Daigneault et al., 2024), or to assess regional forest carbon levels under alternative management scenarios with potential impacts on forest policy (Brown et al., 2018; Boisvenue et al., 2022).

Despite the potential of FDMs to inform forest management practices to promote ecosystem resilience and adaptation, and enhance long-term carbon stocks, there remains high uncertainty in demographic responses to novel climate and disturbance regimes driving forest dynamics under global change (McDowell et al., 2020). While the ecological complexity and processes represented in FDMs has increased in recent years, there remains large variability in predictions of forest dynamics under novel future conditions (Bugmann and Seidl, 2022). This variability reflects collective uncertainty about both the nature of changing conditions (climate, disturbance) and their effects on forest demographic processes with potentially large impacts on the predicted outcomes of adaptive management (Bugmann et al., 2019; Maréchaux et al., 2021).

Predictions of forest dynamics and associated growth and yield would ideally reflect the different sources of uncertainty affecting their values. This is particularly important in the context of predicting adaptive management outcomes under global change (Radke et al., 2020). Specific sources of uncertainty affecting predictions of forest dynamics include imperfect approximation of forest demographic processes and their responses to novel conditions (model or process uncertainty), uncertainty in the model parameters used to approximate forest dynamics (parameter uncertainty), and variability in forest observations used to inform FDMs (data uncertainty; Van Oijen et al., 2005). Additional uncertainty stems from the potential impacts of unknown future conditions or events (e.g., weather extremes, novel disturbances). Quantifying and accounting for these different sources of uncertainty in predictions of forest dynamics is integral to identifying effective management strategies to conserve forest ecosystems and sustain the diversity of services they provide in the face of changing conditions.

While the importance of accounting for uncertainty when modeling forest dynamics is well understood (Purves and Pacala, 2008), the structure and complexity of contemporary FDMs (often comprising large sets of dependent non-linear functions) make it challenging to quantify and propagate different sources of uncertainty to predictions of growth and yield (Wilson et al., 2019; Itter and Finley, 2024). Under current modeling approaches, FDMs are commonly parameterized using regional forest inventory data separately (independently) for each species and demographic process (Hartig et al., 2012; Van Oijen, 2017). Parameter estimates (e.g., restricted maximum likelihood estimates) are then plugged into the target FDM to generate predictions of forest dynamics. This so called “plug in” approach (Kennedy and O’Hagan, 2001) yields single point predictions of future forest states without direct estimates of uncertainty (Van Oijen, 2017). The alternative is to apply FDMs that provide probabilistic predictions of growth and yield reflecting the full probability distribution of future forest states under simulated management approaches (Wilson et al., 2019; Itter and Finley, 2024).

Few FDMs apply a probabilistic modeling approach because it is computationally challenging often requiring that models are run thousands of times within broader simulation routines (Raiho et al., 2021). When probabilistic approaches are applied, they almost exclusively focus on uncertainty in model parameters alone and do not quantify potential uncertainty in the processes that underlie forest dynamics (e.g., disturbance and demographic responses) and noisy (variable) observations of their outcomes (Purves et al., 2008; Fisher et al., 2010; Wilson et al., 2019; Myllymäki et al., 2024). Recent research has focused on addressing the challenge of generating probabilistic predictions of forest dynamics relying on advancements in spatio-temporal statistical methods. Approaches include the development of statistical emulators that approximate the outcomes of and dependencies within complex FDMs (Raiho et al., 2021), and the development of dynamical statistical models motivated by theoretical models of ecological dynamics (Itter and Finley, 2024). The goal of these approaches is to model the process of forest dynamics in a simple, scalable manner allowing for the integration of FDMs within broader probabilistic frameworks. To this end, a Bayesian hierarchical approach is commonly applied to simplify complex forest dynamics through sensible conditioning, quantify and partition different sources of uncertainty, and synthesize multifarious forest data. These frameworks explicitly quantify uncertainty associated with the underlying ecological process, model parameters, and data, and propagate it to model predictions (Laubmeier et al., 2020).

An important benefit of a dynamical statistical approach is that it allows predictions of forest dynamics to learn from and be refined by new or future data (Wikle and Hooten, 2010). This is important when modeling forest dynamics under global change given that future observations of forest responses to novel conditions may be more informative than the historic observations commonly used to parameterize FDMs (McDowell et al., 2020). From an adaptive management perspective, dynamical statistical models support learning from experiments such as the Adaptive Silviculture for Climate Change project (Nagel et al., 2017), reducing uncertainty in predictions based on direct observations of adaptive management outcomes. Dynamical statistical models further allow for the application of methods to identify and reduce the primary sources of uncertainty affecting predictions of forest dynamics under global change through targeted model refinement, experimentation, and new data collection (Dietze et al., 2018). Despite the potential of dynamical statistical models to quantify uncertainty and propagate it to predictions of forest dynamics, the lack of computationally-tractable FDMs that simulate management on an appropriate spatio-temporal scale in terms implementable by forest managers has limited the application of these frameworks to inform contemporary forest management including management for climate resilience and carbon.

In the current study, we build upon a recently-developed dynamical statistical model (Itter and Finley, 2024) to predict forest growth and yield under alternative management approaches based on continuous forest inventory (CFI) data. The model provides probabilistic predictions of forest growth and yield under alternative management scenarios reflecting uncertainty in the demographic processes underlying forest dynamics, the model parameters used to approximate these processes, and the CFI data used to inform the model. Forest dynamics are approximated at a stand scale by modeling changes in the diameter distribution of individual species over time.

The stand scale approach is important both from a computational perspective, but also because it represents forest dynamics on the scale at which management decisions are made (O’Hara and Nagel, 2013). Alternative management strategies are naturally expressed in terms of target residual stand density by species and diameter class (stems and/or basal area per hectare). The model is applied to predict multi-decadal live aboveground biomass (lAGB) growth and yield under five alternative management strategies using long-term CFI data from the USDA Forest Service Penobscot Experimental Forest.

## Materials and methods

### Penobscot Experimental Forest Data

The Penobscot Experimental Forest (PEF) is a 1600 ha experimental forest in central Maine, USA (44^°^52’N, 68^°^38’ W; Fig. 1). It is jointly managed by the USDA Forest Service and the University of Maine to support long-term applied forest ecology research (Brissette and Kenefic, 2014). The PEF sits within the Acadian forest, a transition zone between mixed hardwood forest to the south and boreal forest to the north, comprising a unique mixture of eastern temperate and boreal species. In the current study, species are pooled into eight representative groups: balsam fir, eastern hemlock, spruce, pine/larch, northern white-cedar, red maple, paper birch, other hardwoods. The natural disturbance regime in the PEF is characterized by small-scale, low-severity disturbances such as windthrow punctuated by periodic eastern spruce budworm (*Choristoneura fumiferana* Clem.) outbreaks causing potentially severe defoliation and mortality of mature balsam fir and to a lessor extent spruce species in the forest (Brissette and Kenefic, 2014).

**Figure 1:**
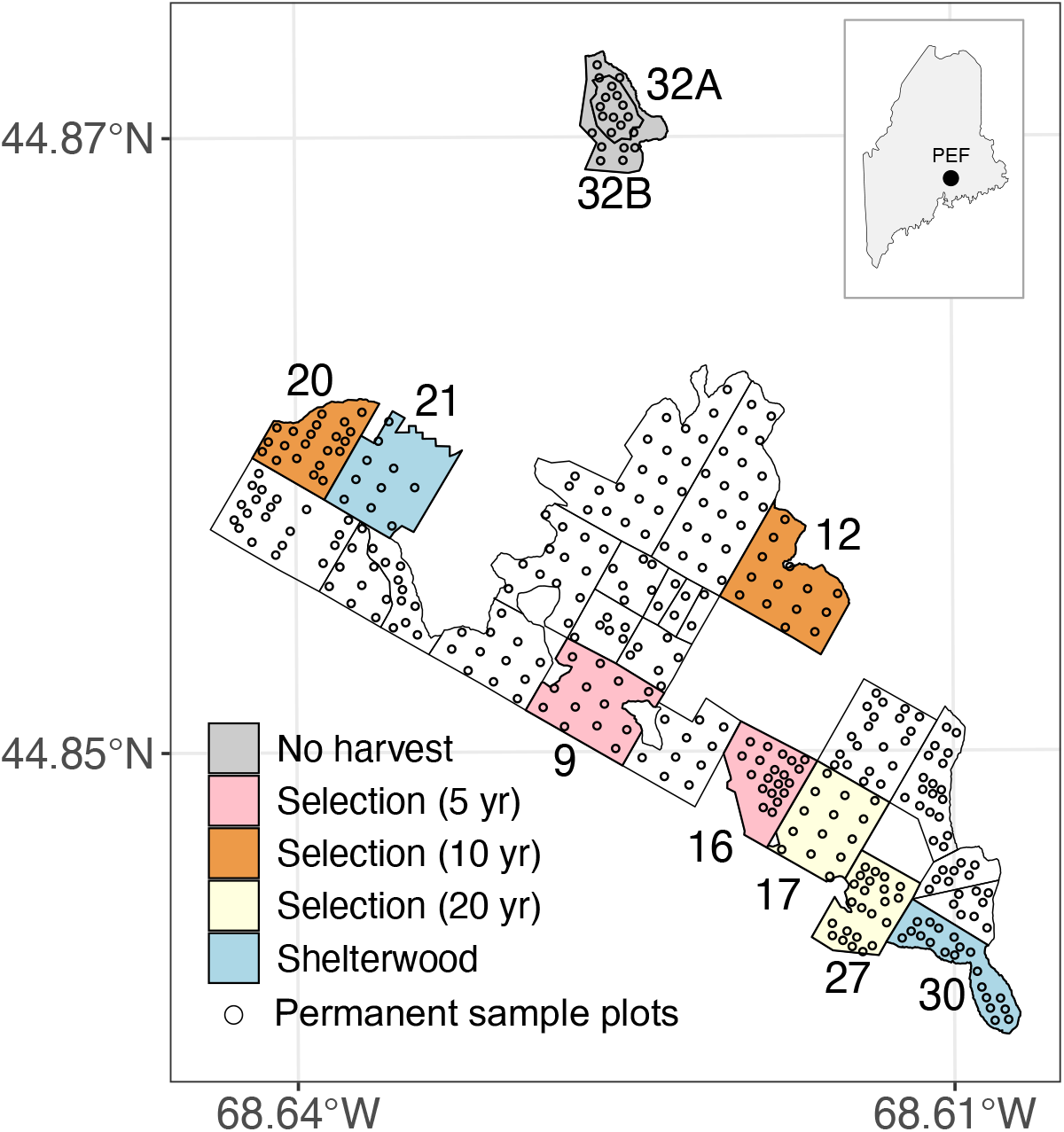
Map of Penobscot Experimental Forest indicating individual management units and location of permanent sample plots. Modeled management units are colored according to silvi-cultural treatment.

A long-term silviculture experiment was established in the PEF between 1952-1957 with replicated even- and uneven-aged silvicultural systems applied across 20 management units (Brissette and Kenefic, 2014). A CFI program was initiated within the PEF shortly after its establishment. Permanent sample plots were installed prior to silvicultural treatments in the 1950s with plots remeasured every 5-10 years through present. Species identity and diameter at breast height (DBH) were recorded for all trees within sample plots using a nested, fixed area design with 0.08 ha plots used to measure large diameter trees (> 11.4 cm) and either 0.02 ha or 0.008 ha nested subplots used to measure small diameter trees (1.3 − 11.4 cm). DBH values were measured to the nearest 0.3 cm (0.1 in) except in historic inventories (prior to 1974) in which tree diameter was classified into 2.5 cm (1.0 in) DBH classes. The number of permanent sample plots measured and specific inventory years vary by management unit (Table 1). With few exceptions, management units were inventoried in the years immediately preceding and following harvest activities providing observations of pre- and post-harvest stand conditions and records of the trees removed in each harvest event (removal tree records begin in 1977).

**Table 1:** Summary of silvicultural treatments and inventory measurements by management unit (MU) within the Penobscot Experimental Forest (harvest year is adjusted to match the first model year post harvest if it occurs in the same year as the pre-harvest inventory).

| MU | Treatment | Model period | Harvest years | Number of inventories | Number of plots |
| --- | --- | --- | --- | --- | --- |
| 32A<br>32B | Control<br>(no harvest) | '75-'14 | NA | 7 | 10 |
| 9<br>16 | Selection system<br>(5-yr cycle) | '78-'14<br>'77-'14 | '84, '89, '94, '99, '09<br>'82, '87, '91, '97, '02, '12 | 13 | 13<br>20 |
| 12<br>16 | Selection system<br>(10-yr cycle) | '75-'14<br>'77-'14 | '84, '95<br>'87, '98 | 9 | 14<br>21 |
| 17<br>27 | Selection system<br>(20-yr cycle) | '76-'14<br>'77-'14 | '95<br>'97 | 8 | 14<br>23 |
| 21<br>30 | Shelterwood +<br>thinning | '76-'14 | '13 | 8 | 10<br>20 |

In the current study, we model forest dynamics using previously published PEF CFI data (Kenefic et al., 2015) for five of the experimental silvicultural systems including three selection systems with varying cutting cycle lengths of 5, 10, and 20 years, a two-stage shelterwood system with one commercial thinning, and a control system under which no harvests were conducted following establishment of the PEF (Table 1). Each silvicultural system was replicated on two management units within the PEF (Fig. 1). Forest dynamics are modeled for approximately 40 years in each of the ten management units (*≈* 1975-2014). The start of the model period varies slightly by management unit as it is adjusted to ensure records of tree removals are available for each harvest event (Table 1).

### Probabilistic growth and yield model

We define a probabilistic forest growth and yield model built around CFI data. The model incorporates management into a recently-developed dynamical statistical model to predict forest dynamics over time (Itter and Finley, 2024). The model is defined for tally data providing observations of the number of individuals by species and DBH class located within permanent sample plots measured periodically as part of a CFI program. We chose to develop the model for tally data instead of continuous DBH observations because such data remains common, particularly in historic inventories and in timber cruises used to support management decisions. Defining a model for tally data allows us to work with a wide range of inventory data since continuous DBH observations can be converted into DBH classes but not vice versa. The dynamical statistical model is defined using a Bayesian hierarchical approach including a data model, process model, and multiple parameter models (Wikle and Hooten, 2010).

#### Data model

We use 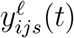 to denote the stem count within DBH class *i* (*i* = 1, …, *k*), from species *j* (*j* = 1, …, *m*), in permanent sample plot *ℓ* (*ℓ* = 1, …, *L*), in inventory year *t* (*t* = 1, …, *T*) within management unit *s* (*s* = 1, …, *S*). We drop management units (*s*) from model notation throughout for simplicity, but note that the dynamical model is fit to all management units simultaneously (i.e., applying a single joint dynamical framework). Plot-level stem counts are modeled using a negative binomial distribution conditional on the latent (unobserved) density of trees per hectare in DBH class *i* from species *j* in year *t* denoted by *λ*_*ij*_(*t*) and a species-specific overdispersion parameter *ϕ*_*j*_ that accounts for variability in stem counts across sampled inventory plots. The data model is defined as

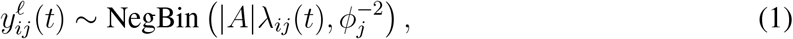

where |*A*| is the size of the permanent sample plot expressed in hectares. In the current analysis, we apply 2.5 cm (1.0 in) DBH classes consistent with the structure of PEF inventories prior to 1974.

#### Process model

The latent stem density *λ*_*ij*_(*t*) describes the true, unobserved state of the modeled forest ecosystem at a given time point. The collection of stem densities for all DBH classes (1 to *k*) and species (1 to *m*) defines the size-species distribution (Clark et al., 2016), which evolves over time driven by forest dynamics. We apply a matrix projection approach to model annual changes in the size-species distribution conditional on a time-dependent propagator matrix **H**_*j*_(*t*) that describes species-specific demographic rates (growth, mortality, regeneration) and forest management (i.e., timber harvesting), and a multiplicative process error term *η*_*ij*_(*t*) that allows for uncertainty in the predicted changes in the latent stem density. The matrix projection process model is defined as

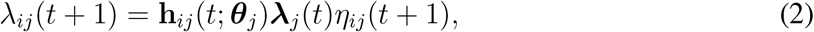

where **h**_*ij*_(*t*; ***θ***_*j*_) is the *i*th row of the (*k × k*) propagator matrix **H**_*j*_(*t*; ***θ***_*j*_) that depends on a set of species-specific demographic parameters ***θ***_*j*_. The ***λ***_*j*_(*t*) term denotes a *k*-dimensional vector of latent DBH class densities for the *j*th species: ***λ***_*j*_(*t*) = (*λ*_1*j*_(*t*), *λ*_2*j*_(*t*), …, *λ*_*kj*_(*t*))^⊺^ (where ^⊺^ indicates the transpose). The propagator matrix **H**_*j*_(*t*; ***θ***_*j*_) describes changes in the latent stem density on an annual basis (*t* corresponds to years). The matrix can be decomposed into sub-matrices reflecting growth (**G**_*j*_), mortality (**M**_*j*_), and forest management intervention (**D**_*j*_) building on common applications of matrix projection models for forest dynamics (Liang and Picard, 2013). The propagator matrix is defined as

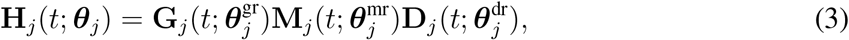

where 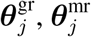 and 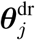 are parameters describing species-specific growth, mortality, and disturbance rates 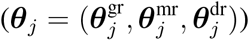. The growth matrix (**G**_*j*_) defines the proportion of individuals in a given DBH class growing into the next larger DBH class (we assume the Usher property holds such that growth is limited to the next size class in a given time step). The mortality matrix (**M**_*j*_) defines the proportion of individuals that survive in a given DBH class. The disturbance matrix (**D**_*j*_) defines the proportion of individuals removed within a given DBH class during a harvest event.

##### Demographic functions

Forest growth and mortality rates are estimated using a generalized linear modeling approach conditional on species-specific demographic parameters. While a wide range of demographic functions exist to predict forest growth and mortality rates (Weiskittel et al., 2011), we model these processes using a logit link function because of its relative simplicity and natural 0-1 constraint. The growth rate of a target DBH class *i* is modeled as a function of DBH and the basal area of all trees in larger DBH classes (so called “basal area larger” abbreviated BAL) both standardized to be on a common scale,

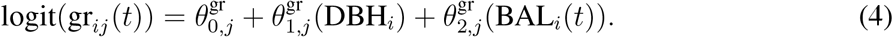

The mortality rate of a target DBH class *i* is comparably modeled as a function of a single parameter representing the species-specific mean mortality rate across all DBH classes,

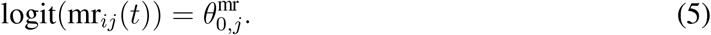

The form of the demographic functions is flexible and supports the inclusion of a wide range of additional predictor variables potentially representing non-linear dynamics. We explored using a range of combinations of DBH and BAL to model growth and mortality rates including quadratic terms for each variable. We further explored modeling mortality as a function of the modeled growth rate given the known dependence between these processes (Cailleret et al., 2017; Camac et al., 2018). The specific variables included here follow from the application of the model to the PEF CFI data considering both the amount of information to estimate regression coefficients and the relative importance of each variable in the model (i.e., whether coefficient estimates were non-zero for most species). The specific variables included here should not be interpreted as explaining growth and mortality on a general level. Rather, we use these specific variables to demonstrate the dynamical model framework and its ability to predict complex forest dynamics in a simple, scalable manner.

Regeneration is notably absent from the definition of the propagator matrix (Eqtn. 3). This can be included through an additive sub-matrix 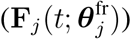 defining species-specific ingrowth of stems into the smallest DBH class (Itter and Finley, 2024). We do not include this regeneration matrix here because there are limited observations of ingrowth in the PEF data (limited to eastern hemlock and balsam fir in a small subset of inventory plots and years). Regeneration is a complex process defined by strong periodicity (intervals with almost no reproductive output punctuated by high seed production years). Without sufficient data it can be challenging to represent the stochastic nature of regeneration and estimate associated model parameters (Díaz-Yáñez et al., 2024). Rather than applying a poorly constrained demographic function, regeneration is approximated in the dynamical model through the species-specific multiplicative process error term for the smallest DBH class (*η*_1*j*_(*t*)). This flexible error term adjusts the latent stem density upward during years when there is observed ingrowth into the smallest DBH class. Application to the PEF data suggests this error term is sufficient to capture stochastic regeneration events without increasing model complexity for sparse observations of ingrowth.

##### Forest management

Modeling the temporal evolution of the size-species distribution allows for straightforward simulation of alternative forest management approaches within the dynamical statistical model. Management interventions are approximated through **D**_*j*_ in Eqtn. (3) with values equal to the residual proportion of stems per hectare remaining following timber removal by species and DBH class. The effect of management within the dynamical model is to reduce the BAL for select species and size classes thereby modifying demographic rates post harvest. That is, the primary impact of management is to modify the competitive growing environment in the stand with corresponding demographic responses following from the relative changes to the size-species distribution.

In order to make meaningful predictions of forest demographic responses to alternative management approaches, the dynamical model must first be trained using inventory observations from past harvest events. There are several ways to approximate the residual stem density following past harvest events. Pre- and post-harvest inventory data often exists for experimental forests where the goal is to understand forest responses to changes in composition, density, size, and structure facilitated by management. When such data exists, it can be used to estimate the proportion of stems per hectare removed across different species and DBH classes. The PEF data used in the current study, for example, provides inventory observations in the years prior to and following experimental management interventions along with records of the individual trees removed during each harvest. The residual stem density is then estimated based on the mean removals across all inventory plots as follows,

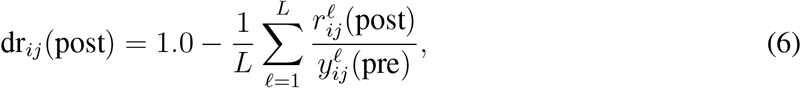

where “post” and “pre” are used to denote inventory years immediately preceding and following a harvest and 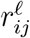 is used to indicate the number of trees removed by species, DBH class, and inventory plot (Fig. S1). The **D**_*j*_(*t*) matrix is fixed at 1.0 in non-harvest years (since no trees were removed) and to the estimated residual stem density (Eqtn. 6) in harvest years. If observations of removal trees do not exist, the residual post-harvest stem density can alternatively be estimated based on pre- and post-harvest inventory data applying flexible modeling approaches such as basis functions, kernel convolutions, or Gaussian processes (Hefley et al., 2017). The same approach can be used to estimate natural disturbance effects on the size-species distribution allowing for prediction of post-disturbance forest dynamics (in this case, **D**_*j*_ represents general forest disturbance). We do not model natural disturbance in the current study because there were have been limited natural disturbances since the PEF was established beyond small scale windthrow events (Granstrom et al., 2022). While these small scale disturbances likely affected the modeled PEF forest dynamics, we do not have sufficient observations of their timing or impacts to incorporate them into the process model. Instead, we rely on the flexible process error term to capture unexplained variation in PEF forest dynamics some of which may be attributable to disturbance.

Simulating forest dynamics under future management scenarios is straightforward under the dynamical statistical model requiring only the specification of a target residual size-species distribution. Once a target residual size-species distribution is set, the entries of the disturbance matrix (**D**_*j*_) are given by,

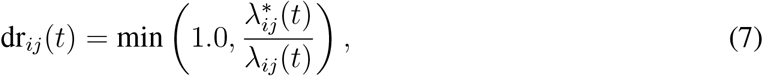

where 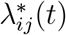 is the target stem density for DBH class *i* and species *j*. Any management intervention is possible under this approach so long as a residual size-species distribution can be determined (Fig. 2). Note that if a forest manager knows the target residual basal area per hectare (rather than stem density) by species and size class, the residual proportion of stems can be determined by transforming the size-species distribution to basal area. The flexibility of the dynamical model to represent management intervention adds to its potential to predict the outcomes of adaptive approaches, which do not fit the definition of classical silvicultural treatments.

**Figure 2:**
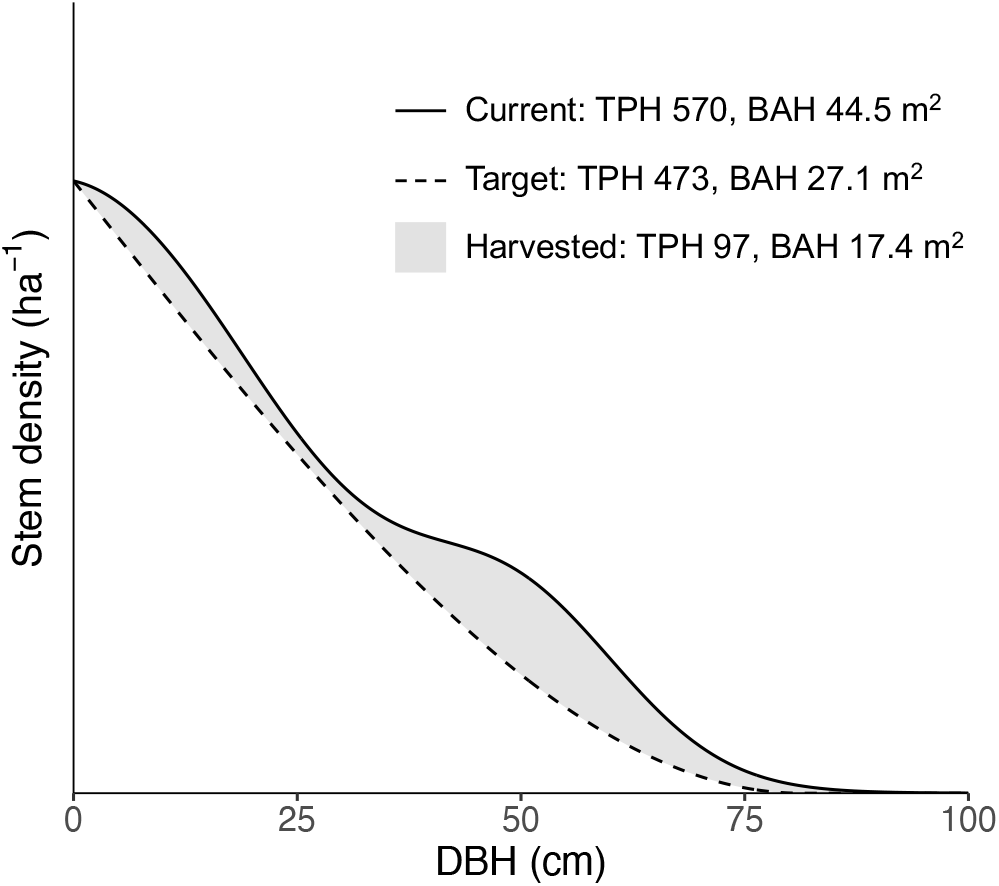
Modeled current stand-level size distribution relative to the desired target distribution. Harvest removals are equal to the difference between the two distributions. Example harvest reduces stand basal area per hectare by 40 percent and moves it from an overstocked, irregular size distribution to a nearly fully stocked, balanced size distribution. Based on thinning depictions provided in Smith et al. (1997).

### Parameter models & implementation

The dynamical statistical model is completed by defining prior distributions for unknown model parameters including demographic parameters, observation and process error terms, and parameters controlling the initial size distribution for all species groups within modeled PEF management units. The model was fit to the PEF CFI data simultaneously (jointly) across all modeled management units using Markov chain Monte Carlo (MCMC) simulation to sample from the joint posterior distribution. We applied a Hamiltonian Monte Carlo MCMC algorithm written in STAN (Stan Development Team, 2024) and implemented using the CmdStanR package (Gabry et al., 2024) for the R statistical computing environment (R Core Team, 2023). Details on parameter models including the specification of prior distributions and model implementation are provided in the Supplementary Material. We further provide all model files and associated data processing scripts (Itter et al., 2025).

### Connecting size-species distribution to forest biomass

Converting the size-species distribution to stand- or population-level variables of interest such as basal area, volume, or biomass is straightforward given applicable allometric functions. Let *z*_*j*_(*t*) denote the species-specific, stand-level value of a forest variable of interest. The value of *z*_*j*_(*t*) is estimated as a function of the size-species distribution as follows,

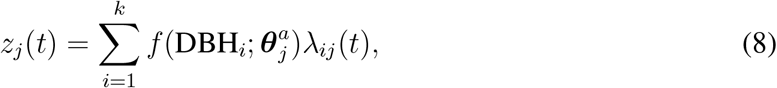

where *f* (DBH_*i*_; 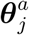) is an allometric function estimating the target variable (basal area, volume, biomass, carbon) as a function of tree DBH given a set of species-specific allometric parameters 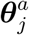. Note the estimation of stand-level variables under the dynamical statistical model closely resembles inventory-based estimates where plot-level values are converted to per hectare values applying the appropriate expansion factor. The size-species distribution is already expressed on a per hectare scale under the dynamical model, so we need only estimate the variable of interest for each species and size class and multiply by the latent stem density (*λ*_*ij*_(*t*)).

In the current study, we convert the size-species distribution into estimates of lAGB per hectare. Live aboveground biomass is estimated applying Jenkins et al. (2003) species-group biomass equations to convert DBH to lAGB (inserted as *f* (DBH_*i*_; 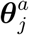) in Eqtn. 8).

## Results

The dynamical model provides probabilistic predictions of the size-species distribution within each management unit in each year of the model period. Changes in the size-species distribution are driven by modeled growth, mortality, and regeneration along with harvest interventions that remove stems from select species and size classes. Summing over the modeled size-species distribution provides predictions of the overall stems per hectare in each year in the model period along with associated uncertainty (Fig. 3). Note that while the minimum size class included in the model is 2.5 cm, predicted stem density dynamics are shown for stems in DBH classes 10.2 cm or larger to avoid depicted dynamics being dominated by large, highly variable small diameter stem densities. The latent overall stem density decreased slowly over time in the two control units (32A, 32B) except during the last 15 years of the model period in unit 32A where stem density increased driven by a large recruitment event of balsam fir in the late 1990s (Fig. 4). While stem density decreased following single tree and group selection harvests in the six selection system units (9, 16, 12, 20, 17, 27), there was a general increase in stem density in the second half of the model period (Fig. 3) due to growth of shade tolerant balsam fir and eastern hemlock saplings into the smallest size class depicted (10.2 cm; Fig. 4). Stem density increased over time in both shelterwood units (21, 30; Fig. 3) driven largely by growth of balsam fir in unit 21, and pine/larch and paper birch in unit 30 (Fig. 4). Stem density decreased at the end of the model period in the two shelterwood units following a commercial thinning in 2013 (Fig. 3).

**Figure 3:**
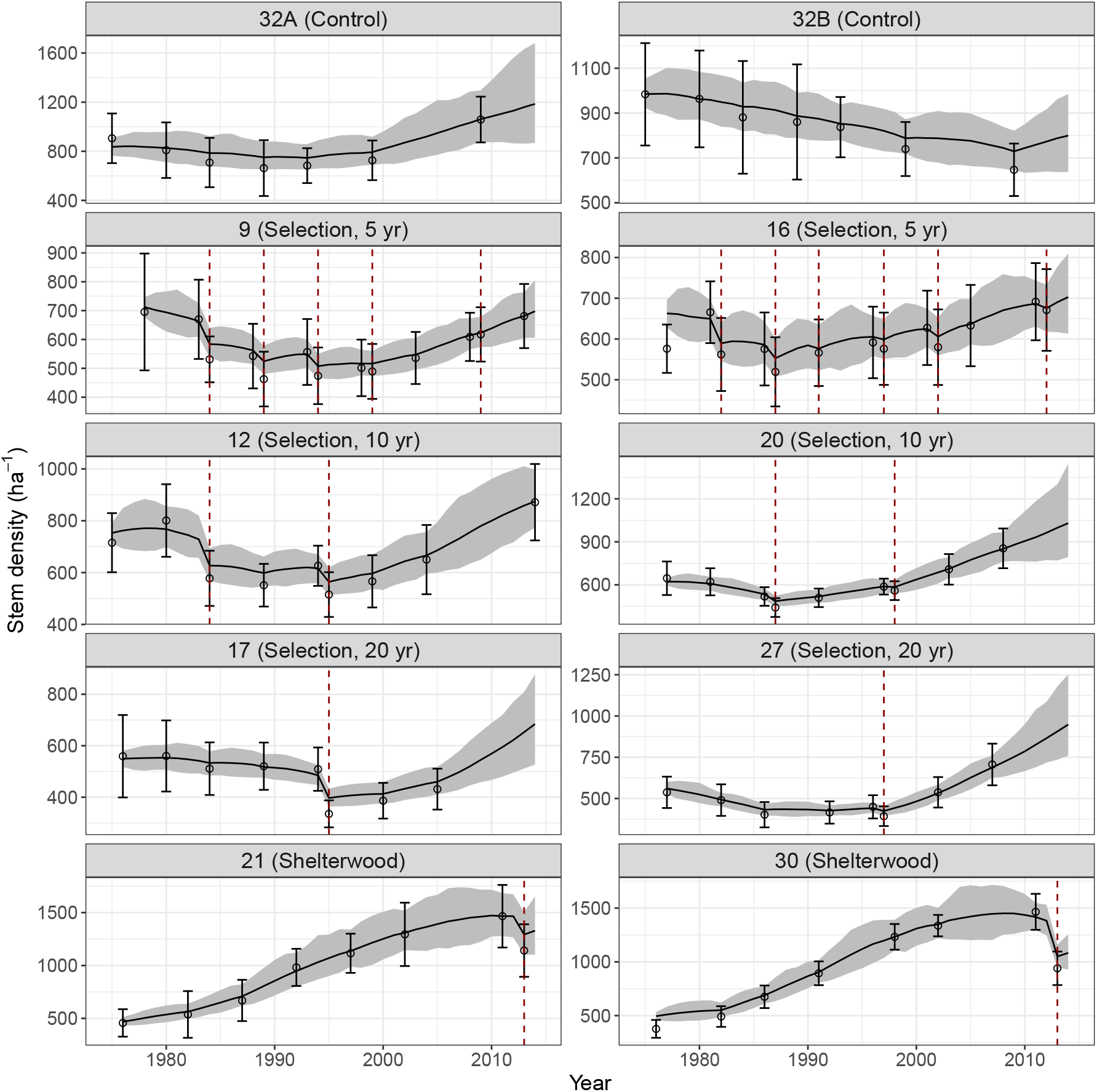
Modeled changes in the overall stem density per hectare within each management unit for trees in DBH classes 10.2 cm or larger. Lines indicate posterior mean density with shaded area indicating the 95 percent credible interval. Points represent the mean overall stem density observed across inventory plots with bars equal to two times the standard error. Red dashed lines indicate harvest years. Treatments and harvest years are defined in Table 1. Note that the range of stem density values varies by management unit.

**Figure 4:**
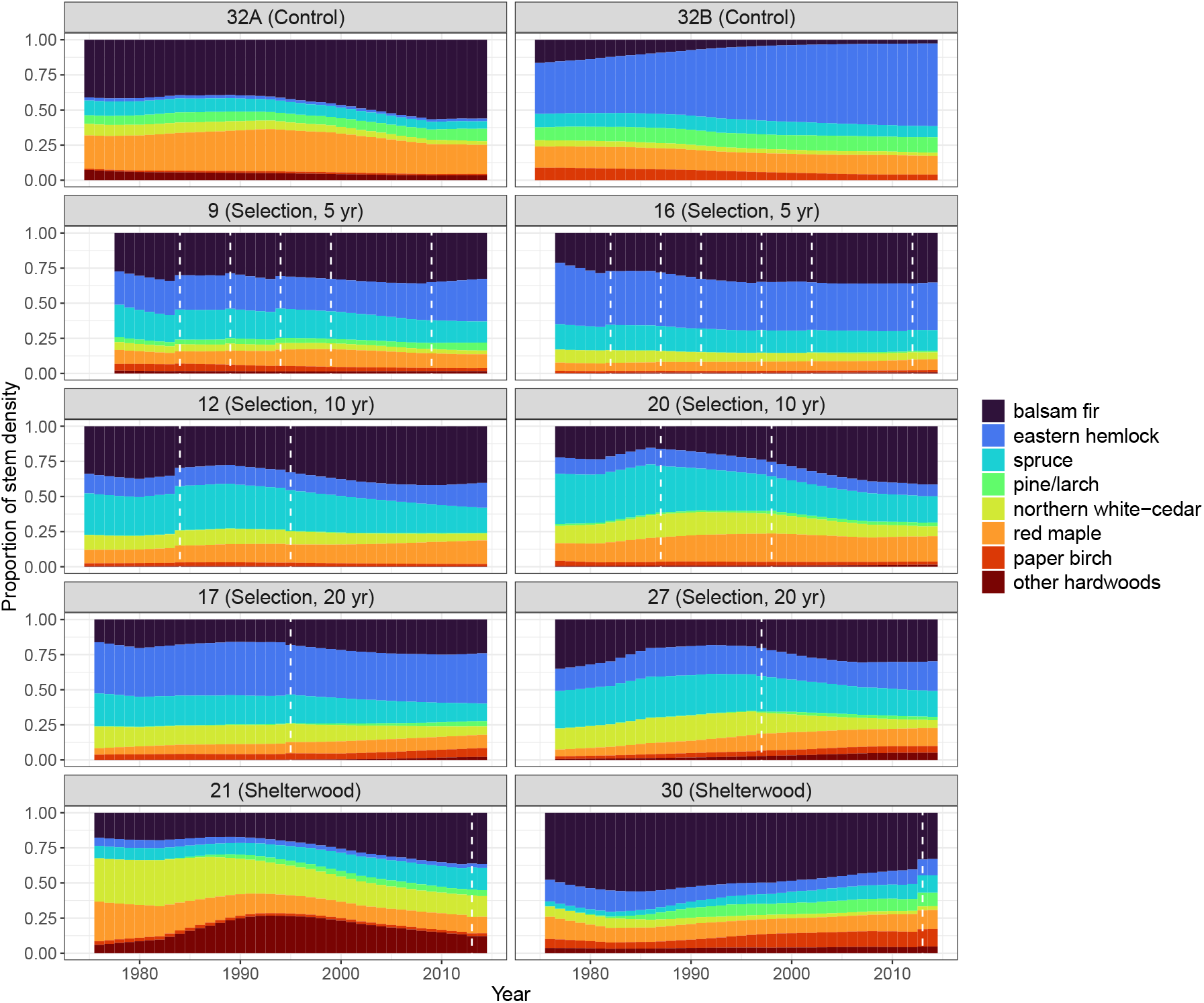
Proportion of overall stem density represented by modeled species groups over time within each management unit for trees in DBH classes 10.2 cm or larger. White dashed lines indicate harvest years. Proportions are based on the posterior mean stem density by species in each year in the model period. Treatments and harvest years are defined in Table 1.

Live aboveground biomass dynamics are presented in Fig. 5 for each management unit along with periodic inventory-based estimates for reference. Live aboveground biomass increased in each management unit over the model period consistent with stands in the stem exclusion to early understory re-initiation phases of development (Fig. 5). Uncertainty in predictions of lAGB decreased when inventory-based estimates were available to inform the dynamical model as seen through narrower credible interval widths in inventory years (Fig. 5). The control units had the highest lAGB over the model period with values between 100-500 tons per hectare, while the shelterwood units had the lowest lAGB with values between 25-200 tons per hectare. Timber harvests resulted in short term reductions in modeled lABG followed by increased lAGB accrual rates post harvest. Balsam fir and eastern hemlock represented large proportions of lAGB in all management units often constituting the majority of lAGB (Fig. 6). Spruce represented a large proportion of lAGB in selection units given its high stem density in these units (Figs. 3, 4). Pine/larch represented a large proportion of lAGB in control units with biomass concentrated in larger DBH, canopy dominant individuals present at lower stem densities (Fig. 4).

**Figure 5:**
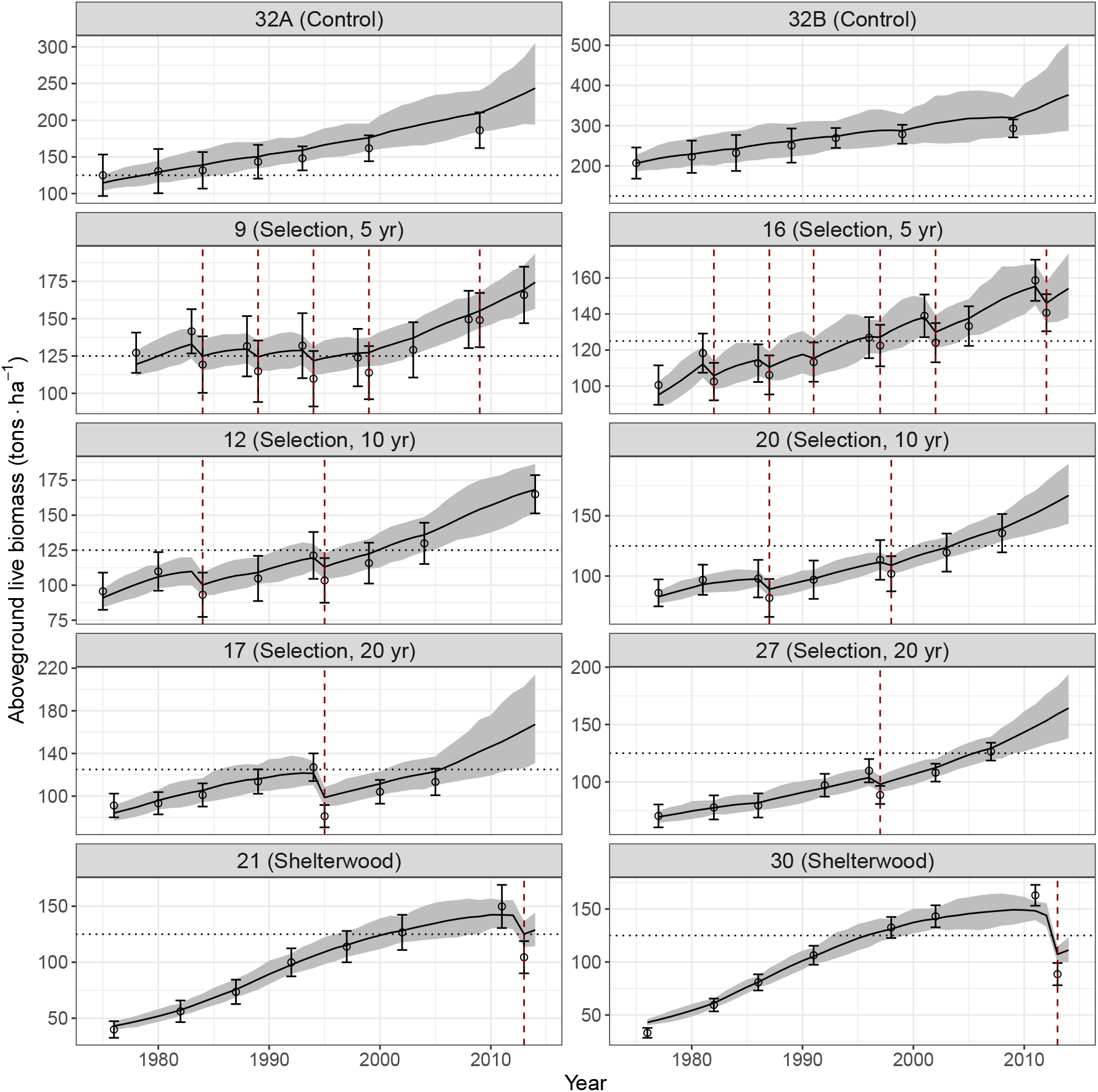
Modeled changes in the live aboveground biomass per hectare within each management unit. Lines indicate posterior mean density with shaded area indicating the 95 percent credible interval. Points represent the mean live aboveground biomass observed across inventory plots with bars equal to two times the standard error. Red dashed lines indicate harvest years. The range of biomass values varies by management unit. A dotted line is provided at 125 tons per ha for reference. Treatments and harvest years are defined in Table 1.

**Figure 6:**
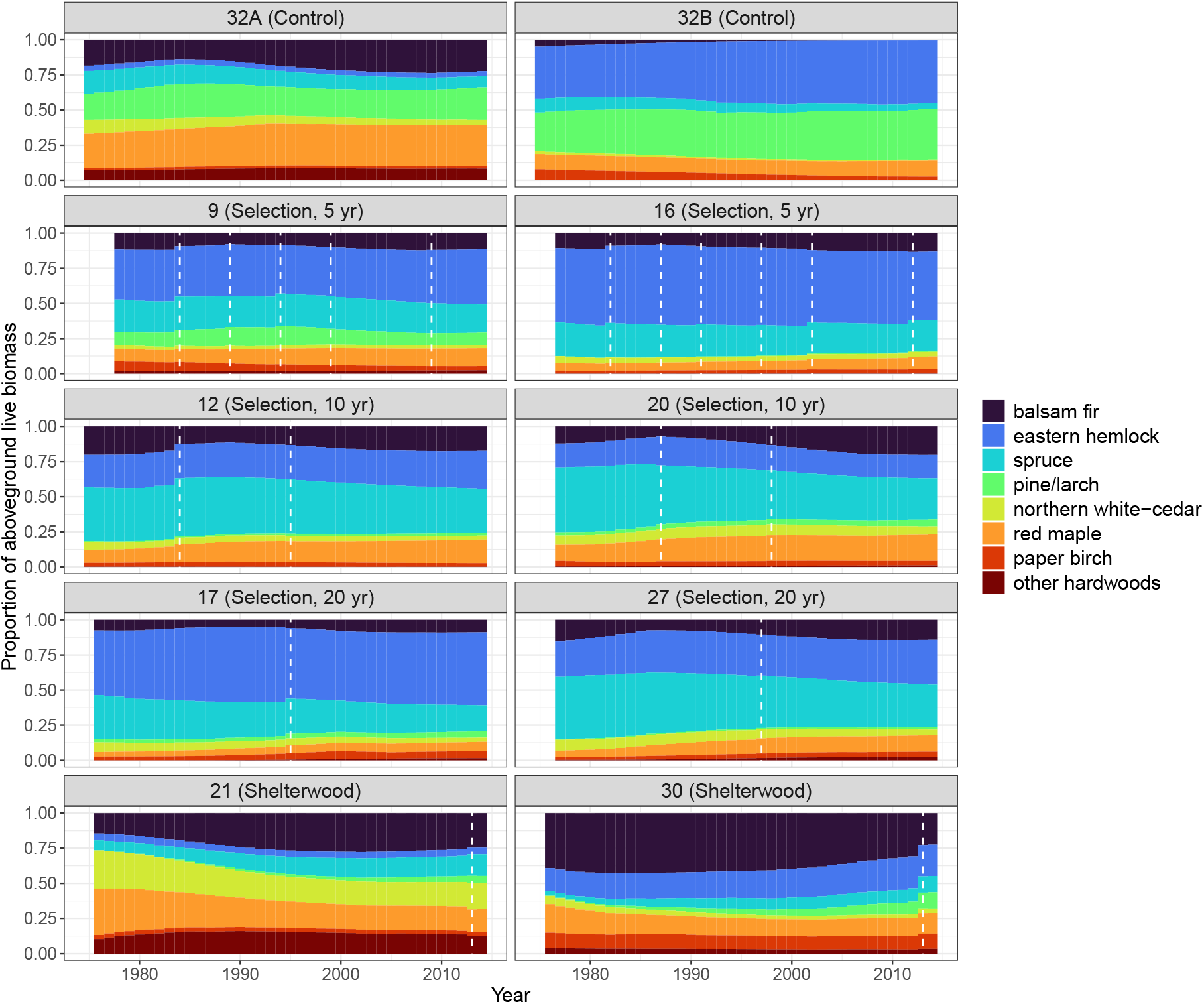
Proportion of live aboveground biomass represented by modeled species groups over time within each management unit. White dashed lines indicate harvest years. Proportions are based on the posterior mean live aboveground biomass by species in each year in the model period. Treatments and harvest years are defined in Table 1.

Predictions of lABG density over time were converted into estimates of growth and yield for select time periods. Specifically, the lAGB dynamics presented in Figure 5 were converted into estimates of periodic annual increment from 1980-2010 and yield in 2010 within each modeled management unit (Table 2). The two control units (32A, 32B) had both the highest lAGB growth and yield over the period of interest. Shelterwood units had the second-highest lAGB growth rates, but the lowest yield on average. Selection units were intermediate in their lAGB growth and yield values. On average, selection units with short cutting cycles (9, 16) had relatively low lAGB growth rates, but higher yields while units with long cutting cycles had higher lAGB growth rates, but lower yields.

**Table 2:**
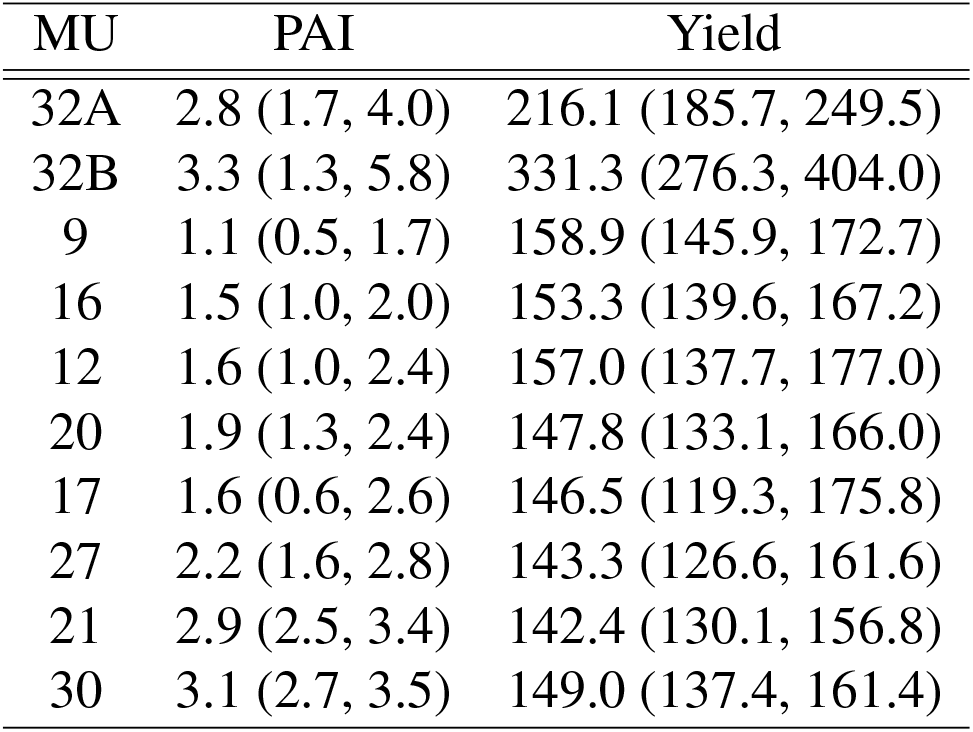
Modeled per-hectare aboveground live biomass periodic annual increment (PAI) between 1980-2010 and yield in 2010 by management unit (MU). Posterior mean values are reported with 95 percent credible intervals in parentheses.

| MU | PAI | Yield |
| --- | --- | --- |
| 32A | 2.8 (1.7, 4.0) | 216.1 (185.7, 249.5) |
| 32B | 3.3 (1.3, 5.8) | 331.3 (276.3, 404.0) |
| 9 | 1.1 (0.5, 1.7) | 158.9 (145.9, 172.7) |
| 16 | 1.5 (1.0, 2.0) | 153.3 (139.6, 167.2) |
| 12 | 1.6 (1.0, 2.4) | 157.0 (137.7, 177.0) |
| 20 | 1.9 (1.3, 2.4) | 147.8 (133.1, 166.0) |
| 17 | 1.6 (0.6, 2.6) | 146.5 (119.3, 175.8) |
| 27 | 2.2 (1.6, 2.8) | 143.3 (126.6, 161.6) |
| 21 | 2.9 (2.5, 3.4) | 142.4 (130.1, 156.8) |
| 30 | 3.1 (2.7, 3.5) | 149.0 (137.4, 161.4) |

Modeled species-specific demographic rates are presented in Table 3. The pine/larch group had both the highest estimated mean growth (0.09) and mortality (0.06) rates. All other species were estimated to have low (*<* 0.05) mean growth and mortality rates highlighting the slow rate of forest change in the modeled management units. The effects of DBH and BAL on modeled growth rates are presented in Table 4. There was limited evidence of an effect of DBH on the annual growth rates of modeled species with the exception of red maple (small effect sizes, 95 percent credible intervals overlap with 1.0). There was evidence of a positive effect of DBH on red maple growth with baseline growth rates increasing slightly (1.1 times on average) with an increase in DBH class. Modeled growth rates were more sensitive to BAL with evidence of a negative effect of BAL on the growth of balsam fir, spruce, pine/larch, and other hardwoods, and a positive effect on the growth of red maple.

**Table 3:** Modeled species-specific growth and mortality rates calculated applying Eqtns. 4 and 5, respectively, conditional on posterior samples of demographic parameters. Growth rate is estimated applying a 10.2 cm diameter at breast height (DBH) class and basal area larger value of 25 m^2^ *·* ha. Values indicate the annual proportion of stems that grow into the next larger DBH class or perish due to mortality. Posterior mean rates are reported with 95 percent credible intervals in parentheses.

| Species | Growth | Mortality |
| --- | --- | --- |
| balsam fir | 0.03 (0.02, 0.04) | 0.02 (0.01, 0.03) |
| eastern hemlock | 0.04 (0.03, 0.04) | 0.01 (0.00, 0.01) |
| spruce | 0.03 (0.02, 0.04) | 0.01 (0.00, 0.02) |
| pine/larch | 0.09 (0.07, 0.10) | 0.06 (0.04, 0.08) |
| northern white-cedar | 0.05 (0.04, 0.07) | 0.02 (0.02, 0.03) |
| red maple | 0.04 (0.03, 0.05) | 0.01 (0.00, 0.02) |
| paper birch | 0.03 (0.02, 0.04) | 0.02 (0.01, 0.03) |
| other hardwoods | 0.03 (0.02, 0.04) | 0.03 (0.01, 0.05) |

**Table 4:** The estimated effects of diameter at breast height (DBH) and basal area larger (BAL) on modeled species-specific growth rates. Values indicate the multiplicative effect of a one unit increase in DBH or BAL on baseline growth as presented in Table 3. Posterior mean rates are reported with 95 percent credible intervals in parentheses. Effect estimates are bolded if the 95 percent credible interval does not overlap with 1.0 (indicative of a measurable effect).

| Species | DBH | BAL |
| --- | --- | --- |
| balsam fir | 0.97 (0.93, 1.02) | **0.97 (0.95, 0.99)** |
| eastern hemlock | 1.03 (0.98, 1.06) | 0.98 (0.96, 1.01) |
| spruce | 0.97 (0.93, 1.01) | **0.96 (0.93, 0.98)** |
| pine/larch | 0.99 (0.97, 1.01) | **0.95 (0.93, 0.98)** |
| northern white-cedar | 1.01 (0.97, 1.06) | 0.99 (0.97, 1.01) |
| red maple | **1.10 (1.06, 1.13)** | **1.02 (1.00, 1.03)** |
| paper birch | 1.02 (0.96, 1.08) | 0.99 (0.95, 1.02) |
| other hardwoods | 1.02 (0.96, 1.08) | **0.94 (0.91, 0.98)** |

## Discussion

Predictions of forest dynamics under alternative management approaches are integral to informing sustainable management of forests under global change. The application of the presented dynamical model to five alternative management regimes within the PEF highlights its flexibility to represent diverse and complex silvicultural treatments by modifying the size-species distribution within forest stands (Figs. 2, S1). Management effects on the composition and overall stem density of modeled stands are depicted in Figs. 3 and 4. Modeled stem density changes were consistent with stands in the late stem exclusion to early understory re-initiation phases of development characterized by decreasing stem densities for the first 20-25 years of the model period, followed by increasing stem density driven by growth of small diameter, shade tolerant eastern hemlock and balsam fir over the last 15-20 years (Fig. 4). While increased stem density is partially due to regeneration observed as ingrowth into the smallest observed (and modeled) size class (2.5 cm), much of it is attributable to the growth of small diameter trees (DBH classes: 2.5, 5.1, 7.6 cm) that were present at the beginning of the model period and subsequently grew into the minimum size class depicted (10.2 cm) in Figs. 3 and 4.

Modeled stem density dynamics were similar between control units (32A, 32B) and selection system units with a short cutting cycle (9, 16) characterized by frequent, low intensity selection harvests. Higher intensity selection harvests within management units with intermediate (12, 20) and long (17, 27) cutting cycles resulted in increased growth of small diameter eastern hemlock and balsam fir leading to increased stem density post-harvest. The increased growth was driven by modeled reductions in BAL post harvest indicative of increased understory light levels. Shelterwood units (21, 30) exhibited increasing stem densities over time driven by high ingrowth rates of balsam fir and other hardwoods following overstory removal in 1967 roughly ten years prior to the start of the model period (Brissette and Kenefic, 2014).

Control units (32A, 32B) had both the highest lAGB periodic annual increment and yield between 1980-2010 (Table 2). Predicted lAGB dynamics in these units highlights a steady increase in biomass over the model period (Fig. 5) consistent with stands in the stem exclusion stage of development characterized by high biomass accrual rates prior to the slowing of stand-level growth rates during the maturation stage. While lAGB growth and yield was similar for the six selection units, those with shorter cutting cycles had lower growth rates and higher yields relative to those with longer cutting cycles where the opposite was true. The differences in lAGB growth and yield under alternative cutting cycles follows from the intensity of selection harvests implemented in these systems. Under the 20-year cycle, higher intensity harvests were implemented, creating growing space to support increased growth, but reducing lAGB stocks. Lower intensity selection harvests under the 5-year cycle resulted in less growing space to support growth, but had less impact on lAGB stocks. Previous studies assessing carbon dynamics in long-term silvicultural experiments applying repeated thinnings at varying intensity identified a similar trade off between increased carbon increment and decreased carbon stocking as thinning intensity increased (D’Amato et al., 2011). Shelterwood units had high lAGB growth rates, but low yields consistent with early stand development stages characterized by high biomass accrual and low biomass stocks.

Modeled growth and mortality rates were relatively low for all species groups in the PEF (Table 3) contributing to limited predicted changes in the stem density and size of modeled management units (Figs. 3, 5). The pine/larch group had both the highest growth and mortality rates of all modeled species groups (Table 3). White pine represented the majority of individuals in this group and was a dominant canopy species in several management units. The high growth rate of the pine/larch group is likely related to its large relative size (Fig. S2) and its predominance in the canopy providing improved access to light. The high mortality rate of the pine/larch group may be due to the coincidence of the model period with the beginning of the understory re-initiation phase triggered in part by the loss of the initial canopy, in this case, co-dominant pine/larch individuals. It may also be caused by unobserved insect damage and disease including white pine weevil and blister rust the effects of which are captured by the process error term within the dynamical model (Munck et al., 2023).

The estimated effects of DBH and BAL on modeled growth rates were minimal across species groups (Table 4). There was evidence of an effect of DBH on red maple growth alone, while BAL was found to have small effects on five of eight species groups (multiplicative changes of less than *<* 10 percent). There was variability in the direction of the mean effect of DBH on species’ growth rates with mean negative effects estimated for balsam fir, spruce, and pine/larch and mean positive effects for all other species groups. While the specific sources of this variability are unclear, we suspect that some of it is attributable to limited intraspecific variability in DBH leading to confounding between tree size and other important growth factors such as crown class (see below for more detail). The mean effect of BAL was consistently negative among species groups with the exception of red maple (Table 4). Despite being consistently negative, the effect of BAL on species’ growth rates was minimal (all values close to 1.0). It is unknown what caused the positive effect of BAL on red maple growth, but it may be related to BAL being confounded with other factors affecting red maple growth.

As noted above, estimating species-specific demographic parameters within the broader dynamical model was challenging given limited variability in growth, mortality, regeneration, and competitive conditions within PEF species groups. The observed DBH range was low for all species groups (Fig. S2) with most species restricted to the smallest size classes likely in suppressed or intermediate crown classes. Notably, only three species groups (eastern hemlock, spruce, pine/larch) had observed stems in size classes above 38.1 cm with the pine/larch group having the largest observed stem sizes. This limited variability did not allow for the complex relationships between size, competition, growth, mortality, and regeneration to be fully explored. For example, initial demographic functions modeled growth rate as a non-linear function of DBH (allowing for the expected concave relationship between growth rate and size) and mortality rate as a function of growth. There was insufficient information to inform higher order, non-linear DBH effects on growth, however, leading to the simple linear function applied here (Eqtn. 4). In addition to limited variability in size and competitive status (as measured by BAL), there was almost no observed ingrowth into the smallest DBH class (a measure of regeneration) over the model period (see Materials and methods). As such, regeneration was modeled through the species- and size-specific process error term in lieu of a stochastic process model predicting the occurrence and intensity of regeneration events for individual species.

Additional inventory data from a wide variety of sites and stand conditions would likely provide the additional variability needed to develop more ecologically meaningful models of demography. Indeed, regional inventory data is regularly used to parameterize FDMs by inferring species-specific demographic rates under heterogeneous conditions for this reason (Didion et al., 2009). Connecting the dynamical model to large-scale CFI networks such as the USDA Forest Service Forest Inventory and Analysis (FIA) program is an active area of research, supporting both more complex models of demography and broader predictions of forest dynamics. Applying such data within a dynamical statistical framework requires extending the modeling approach presented here, which predicts forest dynamics within a single forest over time, to a spatio-temporal model that predicts forest dynamics across a regional collection of forest sites. The motivation behind a dynamical spatio-temporal model is the prediction of site-level forest responses to adaptive management under future climate and disturbance scenarios which quantify and reflect the uncertainty in underlying data, model, and parameters. Open areas of research related to developing a dynamical spatio-temporal model of forest dynamics informed by large-scale CFI data include: incorporating climate and natural disturbance as drivers of forest dynamics within the process model; extending the model framework to allow for space-varying demographic parameters supporting local model calibration; accommodating high dimensional forest inventory data comprising thousands of plots and hundreds of species. Integrating natural disturbance as a stochastic process driving forest dynamics across space and time is particularly important if the model is to predict adaptive management under global change. The current study provides a proof-of-concept of the dynamical modeling approach and its ability to predict complex forest dynamics over time based on simple demographic functions while work on a spatio-temporal dynamical model continues.

The dynamical modeling approach improves the connection between FDM prediction and adaptive management decisions in several important ways. First, the dynamical model can be integrated within a broader Bayesian decision theoretic framework that quantifies the predicted effectiveness of adaptive management strategies to meet objectives (e.g., promotion of ecosystem resistance and resilience) while explicitly accounting for uncertainty (Dorazio and Johnson, 2003). While qualitative adaptive management decision support frameworks exist (Swanston et al., 2016), the Bayesian decision theoretic framework is unique in that it formally integrates uncertainty into the decision making process based on the posterior predictive distribution of forest outcomes under alternative management approaches (Williams and Hooten, 2016). Second, the dynamical model connects to modern approaches to ecological forecasting that aim to reduce uncertainty in predictions through an iterative process of prediction, experimental manipulation and monitoring, model refinement, and data assimilation (Dietze et al., 2018).

Methodologies that improve the connection between models and data to reduce uncertainty in predictions of forest dynamics and better inform adaptive management are particularly important in the context of forest carbon. In particular, the dynamical model presented here can be used to predict forest carbon storage and sequestration rates under alternative management practices with explicit uncertainty (Fig. 5, Table 2). These estimates can be used to quantify the ecological and economic costs and benefits of Improved Forest Management or “Climate-Smart Forestry” practices (Cooper and MacFarlane, 2023). Further, the dynamical model provides a mechanism to assimilate future observations of forest carbon outcomes resulting from such practices supporting the refinement of both forest carbon models and management. Finally, if the dynamical model is integrated within a Bayesian decision theoretic framework, it can be used to account for uncertainty in forest carbon outcomes when deciding on the specific Improved Forest Management practices in which to invest to maximize carbon stores over time.

The dynamical model presented here represents an important first step towards improved quantification of uncertainty in predictions of forest responses to adaptive management. The model generates a probability distribution of predicted forest conditions reflecting uncertainty in the underlying FDM, the parameters upon which the model depends, and the noisy forest inventory observations used to inform it. Ongoing work is focused on connecting the dynamical model to large-scale CFI networks (e.g., USDA Forest Service FIA) to predict forest dynamics over space and time as a function of climate and natural disturbance. The goal of this work is to better inform the sustainable management and conservation of forest ecosystems under global change.

## Conflict of interest statement

The authors declare that they have no conflict of interest.

## Acknowledgments

This study was supported by funding from the National Science Foundation (DEB 1946007) and the USDA Forest Service (23-JV-11242305-080). This work further benefited from valuable discussions among members of the USDA Forest Service Forest Carbon Modeling Group.

## Notes

### Competing Interest Statement

The authors have declared no competing interest.

### Summary of Updates

Integrate small content changes to the manuscript text and figures

## References

Ameray, A., Bergeron, Y., Valeria, O., Montoro Girona, M., and Cavard, X. (2021). Forest carbon management: A review of silvicultural practices and management strategies across boreal, temperate and tropical forests. Current Forestry Reports, pages 1–22.

Boisvenue, C., Paradis, G., Eddy, I. M., McIntire, E. J., and Chubaty, A. M. (2022). Managing forest carbon and landscape capacities. Environmental Research Letters, 17(11):114013.

Brissette, J. C. and Kenefic, L. S. (2014). Centerpiece of research on the Penobscot Experimental Forest: The US Forest Service long-term silvicultural study. In Penobscot Experimental Forest: 60 years of research and demonstration in Maine, 1950–2010, pages 31–57. USDA Forest Service, Gen. Tech. Rep. NRS-P-123, Northern Research Station.

Brown, M. L., Canham, C. D., Murphy, L., and Donovan, T. M. (2018). Timber harvest as the predominant disturbance regime in northeastern us forests: effects of harvest intensification. Ecosphere, 9(3):e02062.

Bugmann, H. and Seidl, R. (2022). The evolution, complexity and diversity of models of long-term forest dynamics. Journal of Ecology, 110(10):2288–2307.

Bugmann, H., Seidl, R., Hartig, F., Bohn, F., Bruna, J., Cailleret, M., François, L., Heinke, J., Henrot, A.-J., Hickler, T., et al. (2019). Tree mortality submodels drive simulated long-term forest dynamics: Assessing 15 models from the stand to global scale. Ecosphere, 10(2):e02616.

Cailleret, M., Jansen, S., Robert, E. M., Desoto, L., Aakala, T., Antos, J. A., Beikircher, B., Bigler, C., Bugmann, H., Caccianiga, M., et al. (2017). A synthesis of radial growth patterns preceding tree mortality. Global change biology, 23(4):1675–1690.

Camac, J. S., Condit, R., FitzJohn, R. G., McCalman, L., Steinberg, D., Westoby, M., Wright, S. J., and Falster, D. S. (2018). Partitioning mortality into growth-dependent and growth-independent hazards across 203 tropical tree species. Proceedings of the National Academy of Sciences, 115(49):12459–12464.

Clark, J. S., Iverson, L., Woodall, C. W., Allen, C. D., Bell, D. M., Bragg, D. C., D’Amato, A. W., Davis, F. W., Hersh, M. H., Ibanez, I., et al. (2016). The impacts of increasing drought on forest dynamics, structure, and biodiversity in the united states. Global Change Biology, 22(7):2329–2352.

Cooper, L. and MacFarlane, D. (2023). Climate-smart forestry: Promise and risks for forests, society, and climate. PLOS Climate, 2(6):e0000212.

Daigneault, A., Simons-Legaard, E., and Weiskittel, A. (2024). Tradeoffs and synergies of optimized management for maximizing carbon sequestration across complex landscapes and diverse ecosystem services. Forest Policy and Economics, 161:103178.

Díaz-Yáñez, O., Käber, Y., Anders, T., Bohn, F., Braziunas, K. H., Bruna, J., Fischer, R., Fischer, S. M., Hetzer, J., Hickler, T., et al. (2024). Tree regeneration in models of forest dynamics: A key priority for further research. Ecosphere, 15(3):e4807.

Didion, M., Kupferschmid, A. D., Lexer, M. J., Rammer, W., Seidl, R., and Bugmann, H. (2009). Potentials and limitations of using large-scale forest inventory data for evaluating forest succession models. Ecological Modelling, 220(2):133–147.

Dietze, M. C., Fox, A., Beck-Johnson, L. M., Betancourt, J. L., Hooten, M. B., Jarnevich, C. S., Keitt, T. H., Kenney, M. A., Laney, C. M., Larsen, L. G., et al. (2018). Iterative near-term ecological forecasting: Needs, opportunities, and challenges. Proceedings of the National Academy of Sciences, 115(7):1424–1432.

Dorazio, R. M. and Johnson, F. A. (2003). Bayesian inference and decision theory—a framework for decision making in natural resource management. Ecological Applications, 13(2):556–563.

D’Amato, A. W., Bradford, J. B., Fraver, S., and Palik, B. J. (2011). Forest management for mitigation and adaptation to climate change: Insights from long-term silviculture experiments. Forest Ecology and Management, 262(5):803–816.

Fisher, R., McDowell, N., Purves, D., Moorcroft, P., Sitch, S., Cox, P., Huntingford, C., Meir, P., and Ian Woodward, F. (2010). Assessing uncertainties in a second-generation dynamic vegetation model caused by ecological scale limitations. New Phytologist, 187(3):666–681.

Gabry, J., Češnovar, R., Johnson, A., and Bronder, S. (2024). cmdstanr: R Interface to ‘Cmd-Stan’. R package version 0.8.1, https://discourse.mc-stan.org.

Giffen, R. A., Ryan, C. M., Belair, E. P., Pounch, M. A., and Brown, S. (2022). Storing more carbon by improving forest management in the Acadian forest of New England, USA. Forests, 13(12):2031.

Granstrom, M., Crandall, M. S., Kenefic, L. S., and Weiskittel, A. R. (2022). Tree quality and value: results in northern conifer stands after 65 years of silviculture and harvest. Canadian Journal of Forest Research, 52(5):794–807.

Griscom, B. W., Adams, J., Ellis, P. W., Houghton, R. A., Lomax, G., Miteva, D. A., Schlesinger, W. H., Shoch, D., Siikamäki, J. V., Smith, P., et al. (2017). Natural climate solutions. Proceedings of the National Academy of Sciences, 114(44):11645–11650.

Gustafson, E. J., Kern, C. C., Miranda, B. R., Sturtevant, B. R., Bronson, D. R., and Kabrick, J. M. (2020). Climate adaptive silviculture strategies: How do they impact growth, yield, diversity and value in forested landscapes? Forest ecology and management, 470:118208.

Hartig, F., Dyke, J., Hickler, T., Higgins, S. I., O’Hara, R. B., Scheiter, S., and Huth, A. (2012). Connecting dynamic vegetation models to data–an inverse perspective. Journal of Biogeography, 39(12):2240–2252.

Hefley, T. J., Broms, K. M., Brost, B. M., Buderman, F. E., Kay, S. L., Scharf, H. R., Tipton, J. R., Williams, P. J., and Hooten, M. B. (2017). The basis function approach for modeling autocorrelation in ecological data. Ecology, 98(3):632–646.

Itter, M. S. and Finley, A. O. (2024). Toward improved uncertainty quantification in predictions of forest dynamics: A dynamical model of forest change. bioRxiv, pages 2024–07.

Itter, M. S., Finley, A. O., and Weiskittel, A. (2025). Model code for ‘Connecting growth and yield models to continuous forest inventory data to better account for uncertainty’. Zenodo. https://zenodo.org/records/14691971?preview=1&token=eyJhbGciOiJIUzUxMiJ9.eyJpZCI6ImYwYjdlOTBiLWMxYWEtNDQ5Ny05ZTM1LTI0ODc2ZjQ1M2JkMSIsImRhdGEiOnt9LCJyYW5kb20iOiI2OWMxZDEzYzIwYjc3NjNjM2NkMTA4Mzc2NWYxNTIxYSJ9.Gy0ucT_vtBKr428SlwT51KvzJblqai9m1K7xXvgp_CLaSkx7bZnza0UJOfmM-tdWgI9Grea0DMHIJt3IU0KD8Q.

Jenkins, J. C., Chojnacky, D. C., Heath, L. S., and Birdsey, R. A. (2003). National-scale biomass estimators for united states tree species. Forest science, 49(1):12–35.

Kenefic, L. S., Rogers, N. S., Puhlick, J. J., Waskiewicz, J. D., and Brissette, J. C. (2015). Overstory tree and regeneration data from the ‘Silvicultural Effects on Composition, Structure, and Growth’ study at Penobscot Experimental Forest. 2nd Edition. Forest Service Research Data Archive, 10.2737/RDS-2012-0008-2.

Kennedy, M. C. and O’Hagan, A. (2001). Bayesian calibration of computer models. Journal of the Royal Statistical Society: Series B (Statistical Methodology), 63(3):425–464.

Laubmeier, A. N., Cazelles, B., Cuddington, K., Erickson, K. D., Fortin, M.-J., Ogle, K., Wikle, C. K., Zhu, K., and Zipkin, E. F. (2020). Ecological dynamics: Integrating empirical, statistical, and analytical methods. Trends in Ecology & Evolution, 35(12):1090–1099.

Liang, J. and Picard, N. (2013). Matrix model of forest dynamics: An overview and outlook. Forest Science, 59(3):359–378.

Maréchaux, I., Langerwisch, F., Huth, A., Bugmann, H., Morin, X., Reyer, C. P., Seidl, R., Collalti, A., Dantas de Paula, M., Fischer, R., et al. (2021). Tackling unresolved questions in forest ecology: The past and future role of simulation models. Ecology and Evolution, 11(9):3746–3770.

McDowell, N. G., Allen, C. D., Anderson-Teixeira, K., Aukema, B. H., Bond-Lamberty, B., Chini, L., Clark, J. S., Dietze, M., Grossiord, C., Hanbury-Brown, A., et al. (2020). Pervasive shifts in forest dynamics in a changing world. Science, 368(6494):eaaz9463.

Mina, M., Bugmann, H., Cordonnier, T., Irauschek, F., Klopcic, M., Pardos, M., and Cailleret, M. (2017). Future ecosystem services from european mountain forests under climate change. Journal of Applied Ecology, 54(2):389–401.

Munck, I. A., Yamasaki, M., and Janelle, J. (2023). Silvicultural treatments improve pest and disease conditions of white pine (pinus strobus) residual trees and regeneration. Frontiers in Forests and Global Change, 6:1239835.

Myllymäki, M., Kuronen, M., Bianchi, S., Pommerening, A., and Mehtätalo, L. (2024). A Bayesian approach to projecting forest dynamics and related uncertainty: An application to continuous cover forests. Ecological Modelling, 491:110669.

Nagel, L. M., Palik, B. J., Battaglia, M. A., D’Amato, A. W., Guldin, J. M., Swanston, C. W., Janowiak, M. K., Powers, M. P., Joyce, L. A., Millar, C. I., et al. (2017). Adaptive silviculture for climate change: a national experiment in manager-scientist partnerships to apply an adaptation framework. Journal of Forestry, 115(3):167–178.

Nunery, J. S. and Keeton, W. S. (2010). Forest carbon storage in the northeastern united states: net effects of harvesting frequency, post-harvest retention, and wood products. Forest Ecology and management, 259(8):1363–1375.

O’Hara, K. L. and Nagel, L. M. (2013). The stand: Revisiting a central concept in forestry. Journal of Forestry, 111(5):335–340.

Pretzsch, H. (2009). Forest dynamics, growth, and yield, volume 684. Springer.

Purves, D. and Pacala, S. (2008). Predictive models of forest dynamics. Science, 320(5882):1452–1453.

Purves, D. W., Lichstein, J. W., Strigul, N., and Pacala, S. W. (2008). Predicting and understanding forest dynamics using a simple tractable model. Proceedings of the National Academy of Sciences, 105(44):17018–17022.

R Core Team (2023). R: A Language and Environment for Statistical Computing. R Foundation for Statistical Computing, Vienna, Austria.

Radke, N., Keller, K., Yousefpour, R., and Hanewinkel, M. (2020). Identifying decision-relevant uncertainties for dynamic adaptive forest management under climate change. Climatic change, 163(2):891–911.

Raiho, A. M., Nicklen, E. F., Foster, A. C., Roland, C. A., and Hooten, M. B. (2021). Bridging implementation gaps to connect large ecological datasets and complex models. Ecology and Evolution, 11(24):18271–18287.

Seidl, R., Albrich, K., Thom, D., and Rammer, W. (2018). Harnessing landscape heterogeneity for managing future disturbance risks in forest ecosystems. Journal of Environmental Management, 209:46–56.

Seidl, R., Rammer, W., Lasch, P., Badeck, F., and Lexer, M. J. (2008). Does conversion of even-aged, secondary coniferous forests affect carbon sequestration? a simulation study under changing environmental conditions. Silva Fennica, 42(3):369.

Smith, D. M., Larson, B. C., Kelty, M. J., Ashton, P. M. S., et al. (1997). The practice of silviculture: applied forest ecology. 9th Ed. John Wiley and Sons, Inc.

Stan Development Team (2024). Stan modeling language users guide and reference manual, 2.34. https://mc-stan.org.

Swanston, C. W., Janowiak, M. K., Brandt, L. A., Butler, P. R., Handler, S. D., Shannon, P. D., Lewis, A. D., Hall, K., Fahey, R. T., Scott, L., et al. (2016). Forest adaptation resources: Climate change tools and approaches for land managers. Gen. Tech. Rep. NRS-GTR-87-2. Newtown Square, PA: US Department of Agriculture, Forest Service, Northern Research Station. 161 p. 10.2737/NRS-GTR-87-2., 87:1–161.

Van Oijen, M. (2017). Bayesian methods for quantifying and reducing uncertainty and error in forest models. Current Forestry Reports, 3:269–280.

Van Oijen, M., Rougier, J., and Smith, R. (2005). Bayesian calibration of process-based forest models: Bridging the gap between models and data. Tree physiology, 25(7):915–927.

Weiskittel, A. R., Hann, D. W., Kershaw Jr, J. A., and Vanclay, J. K. (2011). Forest growth and yield modeling. John Wiley & Sons.

Wikle, C. K. and Hooten, M. B. (2010). A general science-based framework for dynamical spatio-temporal models. Test, 19(3):417–451.

Williams, P. J. and Hooten, M. B. (2016). Combining statistical inference and decisions in ecology. Ecological Applications, 26(6):1930–1942.

Wilson, D., Monleon, V., and Weiskittel, A. (2019). Quantification and incorporation of uncertainty in forest growth and yield projections using a Bayesian probabilistic framework: (A demonstration for plantation coastal Douglas-fir in the Pacific Northwest, USA). Mathematical & Computational Forestry & Natural Resource Sciences, 11(2).

