## Supporting information for "Connecting growth and yield models to continuous forest inventory data to better account for uncertainty"

### S1 Supplemental figures

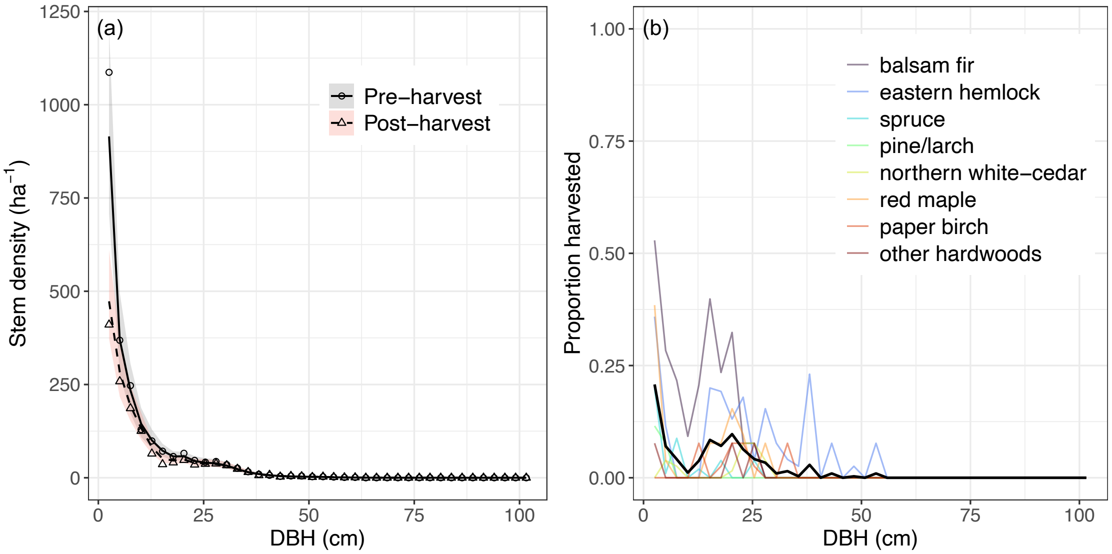


Figure S1: Modeled size distribution of management unit 9 in the years immediately preceding and following a single tree and small group selection harvest in 1984 (a) along with post-harvest inventory estimates of the proportion of each size class removed by species and overall (b). In (a) lines are the posterior mean stem density with shaded areas indicating 95 percent credible intervals and points depicting inventory observations. In (b) species-specific harvest removals are indicated with colored lines while the overall mean is shown in black.

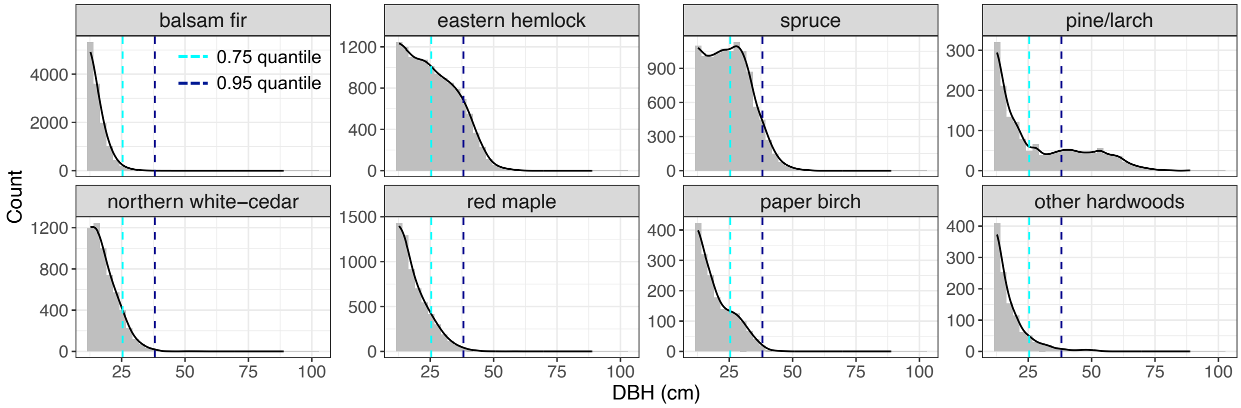


Figure S2: Distribution of observed stem counts by species and size class pooling observations across all management units, permanent sample plots, and inventory years. Minimum size class depicted is 12.70 cm, above which all size classes were measured using a consistent plot size. Quantile values shown are for the overall count data pooled across species (0.75 = 25.4 cm; 0.95 = 38.1 cm).

#### S2 Supplemental modeling information

##### S2.1 Parameter models

The dynamical statistical model is completed by defining distributions for unknown model parameters including demographic parameters, observation and process error terms, and parameters controlling the initial size-species distribution in modeled management units. We assign semi-informative normal prior distributions to each of the demographic parameters,

$$\theta_{\ell,j} \sim \text{Norm}(\mu_\ell, \tau_l^2),$$

with mean values ($\mu_\ell$) set to -3.0, 0.0, and -3.0 for the three growth rate parameters corresponding to an intercept, and the effect of DBH and basal area larger, respectively, and 3.0 for the species-specific mortality rate. All demographic parameters were assigned a prior standard deviation ($\tau_l$) of 5.0. While it is common to apply an exchangeable prior for species-specific demographic parameters allowing for partial pooling of information among species (Itter and Finley, 2024), we choose not to do so here given the small number of species groups (eight) and low variability in their respective demographic rates.

Integral to the dynamical statistical model is the inclusion of process and observation error terms. Process error terms are assigned a log-normal prior,

$$\eta_{ij}(t) \sim \text{LogNorm}(0, \sigma_j^2),$$

where $\sigma_j^2$ is a species-specific, multiplicative process variance parameter. The negative binomial data model (Eqtn. 1) is equivalent to a Poisson data model with a multiplicative observation error term with a Gamma prior with mean 1.0 and variance $\phi_j^2$ (Lawless, 1987). We assign zero-truncated normal priors to the process error ($\sigma_j$) and observation error ($\phi_j$) standard deviation parameters for each species:

$$\phi_j \sim \text{N}_{[0,\infty]}(0, 0.5),$$
$$\sigma_j \sim \text{N}_{[0,\infty]}(0, 0.5).$$

The process error terms are sampled using a non-centered parameterization,

$$\eta_{ij}(t) = \exp(z_{ij}^{\eta}(t)\, \sigma_j),$$

where $z_{ij}^{\eta}(t)$ is assigned an independent standard normal prior.

Initial size-species density values correspond to the latent stem density at the initial time point ($\boldsymbol{\lambda}_j(0)$). We model initial size-species density values using a standardized normal density to represent the size distribution of each species scaled by an estimate of the overall density of the species at the initial time point. Specifically,

$$\lambda_{ij}(0) = n_j f(z_{ij}),$$

where $n_j$ is the initial overall density of species $j$ in terms of trees per hectare and $f(z_{ij})$ is the standardized size density. We apply a standard normal distribution to model the initial size density,

$$f(z_{ij}) = \frac{\phi(z_{ij})}{\sum_{i=1}^{k} \phi(z_{ij})},$$

where $\phi(z_{ij})$ is the standard normal density of $z_{ij}$, which is defined as $z_{ij} = v_j^{-1}(d_i - q_j)$ where $d_i$ is the midpoint of the $i$th size class, $q_j$ is the mean of the initial size distribution, and $v_j$ is its standard deviation. Note that the standardized distribution sums to one across the modeled size classes: $\sum_{i=1}^{k} f(z_{ij}) = 1.0$ for $j = 1, \ldots, m$. We apply a non-centered parameterization to estimate the initial stem density ($n_j$) and mean of the size distribution ($q_j$),

$$\begin{aligned} n_j &= \exp(\tilde{n}_j + z_j^n \, \tau_j^n), \\ q_j &= \tilde{q}_j + z_j^q \, \tau_j^q, \end{aligned}$$

where $\tilde{n}_j$ and $\tilde{q}_j$ are prior estimates of the initial values and $\tau_j^n$ and $\tau_j^q$ are prior uncertainty estimates based on the first inventory observation. Note that the mean of the standard normal distribution used to approximate the initial size distribution may be negative as is likely to be the case for strongly right-skewed size distributions. We assign $z_j^n$ and $z_j^q$ independent standard normal priors for each species. Lastly, we assign a half Cauchy prior to the standard deviation of the initial size distribution: $v_j \sim \text{HalfCauchy}(0, 5)$.

##### S2.2 Model implementation

The dynamical model was fit jointly to all modeled Penobscot Experimental Forest (PEF) management units simultaneously using Markov chain Monte Carlo (MCMC) simulation to sample from the joint posterior distribution. We applied a Hamiltonian Monte Carlo MCMC algorithm written in STAN (Stan Development Team, 2024) and implemented using the CmdStanR package (Gabry et al., 2024) for the R statistical computing environment (R Core Team, 2023). Three MCMC chains were run for a total of 2,000 iterations with a burn-in of 1,000 iterations. Model convergence was assessed based on R-hat values ($< 1.05$), bulk effective sample size (minimum of 100 per chain), and visual inspection of chains for all parameters (Vehtari et al., 2021).

Starting values for demographic rate parameters are generated based on generalized linear growth and mortality models fit to individual tree demographic estimates based on PEF inventory observations. Starting values for process and observation error components as well as the standard deviation of the initial size distribution are sampled from assigned prior distributions. Lastly, starting values for the initial value parameters are generated by fitting the initial size distribution model to the first inventory observation.

### References


Gabry, J., Češnovar, R., Johnson, A., and Bronder, S. (2024). *cmdstanr: R Interface to 'CmdStan'*. R package version 0.8.1, https://discourse.mc-stan.org.

Itter, M. S. and Finley, A. O. (2024). Toward improved uncertainty quantification in predictions of forest dynamics: A dynamical model of forest change. *bioRxiv*, pages 2024–07.

Lawless, J. F. (1987). Negative binomial and mixed Poisson regression. *The Canadian Journal of Statistics/La Revue Canadienne de Statistique*, pages 209–225.

R Core Team (2023). *R: A Language and Environment for Statistical Computing*. R Foundation for Statistical Computing, Vienna, Austria.

Stan Development Team (2024). Stan modeling language users guide and reference manual, 2.34. `https://mc-stan.org`.

Vehtari, A., Gelman, A., Simpson, D., Carpenter, B., and Bürkner, P.-C. (2021). Rank-normalization, folding, and localization: An improved $\widehat{R}$ for assessing convergence of MCMC (with discussion). *Bayesian Analysis*, 16(2):667–718.
